# Propofol anesthesia alters cortical traveling waves

**DOI:** 10.1101/2022.01.11.475912

**Authors:** Sayak Bhattacharya, Jacob A. Donoghue, Meredith Mahnke, Scott L. Brincat, Emery N. Brown, Earl K. Miller

## Abstract

Oscillatory dynamics in cortex seem to organize into traveling waves that serve a variety of functions. Recent studies show that propofol, a widely used anesthetic, dramatically alters cortical oscillations by increasing slow-delta oscillatory power and coherence. It is not known how this affects traveling waves. We compared traveling waves across the cortex of non-human primates (NHPs) before, during, and after propofol-induced loss-of-consciousness (LOC). After LOC, traveling waves in the slow-delta (∼ 1Hz) range increased, grew more organized, and travelled in different directions relative to the awake state. Higher frequency (8-30 Hz) traveling waves, by contrast, decreased, lost structure, and switched to directions where the slow-delta waves were less frequent. The results suggest that LOC may be due, in part, to changes in slow-delta traveling waves that, in turn, alter and disrupt traveling waves in the higher frequencies associated with cognition.

## Introduction

Traveling waves are spatially organized patterns of activity whose peaks and troughs move sequentially across the brain. They have been observed in a variety of brain areas, including the cortex, and across a wide range of frequencies (from 1 - ∼40 Hz) (Muller et al. 2018; 2014; Takahashi et al. 2011; Muller and Destexhe 2012). Traveling waves were first observed under anesthesia in the visual cortex (Ebersole and Kaplan 1981; Cowey 1964), and later in the auditory (Reimer et al. 2011) and somatosensory cortices (Ferezou, Bolea, and Petersen 2006). They seem abundant under anesthesia (Liang et al. 2021; Townsend and Gong 2018; Nauhaus et al. 2009; Sato, Nauhaus, and Carandini 2012; Benucci, Frazor, and Carandini 2007), perhaps because of lower background noise (Muller et al. 2018). They have also been documented during sleep (Muller et al. 2016; Massimini et al. 2004) and early development (Watt et al. 2009; Wong, Meister, and Shatz 1993). However, there are growing observations of traveling waves in the awake cortex (Takahashi et al. 2011; Sreekumar et al. 2020; Alamia and VanRullen 2019) (and hippocampus (Lubenov and Siapas 2009; Honghui Zhang et al. 2018)). There is also a growing realization of their usefulness and functional relevance. They can, for example, retain recent history of network activations, keep track of time, and may even perform computation (Muller et al. 2018; Muller and Destexhe 2012; Heitmann and Ermentrout 2020; Ermentrout and Kleinfeld 2001). Experimental observations show that they change with task demands and impact behavior (Alamia and VanRullen 2019; Bhattacharya et al. 2021). For example, when traveling waves in visual cortex are better organized, animals are better at detecting targets (Davis et al. 2020).

Traveling waves in both awake and anesthetized animals show specific properties. They can be planar or rotational (Muller et al. 2016; Bhattacharya et al. 2021). They tend to flow in certain directions and have different degrees of organization and scale (Muller et al. 2018). However, they have never been directly compared between anesthetized and awake states. This is particularly relevant because of recent studies showing that propofol, a widely used anesthetic, profoundly affects oscillatory dynamics in cortex (Bastos et al. 2021; Lewis et al. 2012; Purdon et al. 2013; 2015; Redinbaugh et al. 2020). Following loss of consciousness (LOC), the cortex of non-human primates (NHPs) shows strong increases in low-frequency slow-delta (∼1 Hz) local field potential (LFP) power and coherence. This reversed when propofol was ceased and the NHPs regained consciousness. In other words, propofol does not merely “turn off” cortex. It changes cortical rhythms. It is thus likely to affect traveling waves as well.

To investigate this, we examined data from our previous study on the effects of propofol on cortical oscillatory power and coherence (Bastos et al. 2021). We used methods previously employed to identify and document traveling waves in the prefrontal cortex during working memory (Bhattacharya et al. 2021). Recordings were made from electrode arrays in three cortical areas: the ventrolateral prefrontal cortex, frontal eye fields and the auditory parabelt cortex of macaque monkeys. The NHPs transitioned from awake to LOC upon propofol administration and then back to awake state, upon propofol cessation. Propofol-induced LOC increased slow-delta traveling waves, strengthened their organization, and caused them to travel in different directions relative to the awake state. There was a corresponding decrease in higher-frequency (8-30 Hz) traveling waves. In fact, the slow-delta traveling waves seemed to “crowd out” high frequency traveling waves, causing them to flow in directions where the slow-delta waves were less frequent.

## Results

Two macaque monkeys were used. Throughout each experimental session, they were exposed to a classical conditioning paradigm. A conditioned stimulus (tone) preceded a puff of air toward the eyes (Fig. 1A). This was used to test their responsiveness to external stimulation (the airpuff). Propofol was administered (see Methods) in two phases (Fig. 1A). First, a higher dose (280-580 mcg/kg/min) of propofol was administered to induce loss-of-consciousness (LOC). LOC was deemed at the timestamp where the subject’s eyes closed and did not reopen for the remainder of the infusion. Soon afterward, the propofol infusion rate was reduced to a lower level (140-320 mcg/kg/min) that could still maintain LOC. After LOC, the animals did not respond to the airpuff. Later, propofol infusion was ceased. After a brief period, there was recovery of consciousness (ROC), i.e. the eyes opened and/or the animals began to respond to the airpuff.

**Figure 1:**
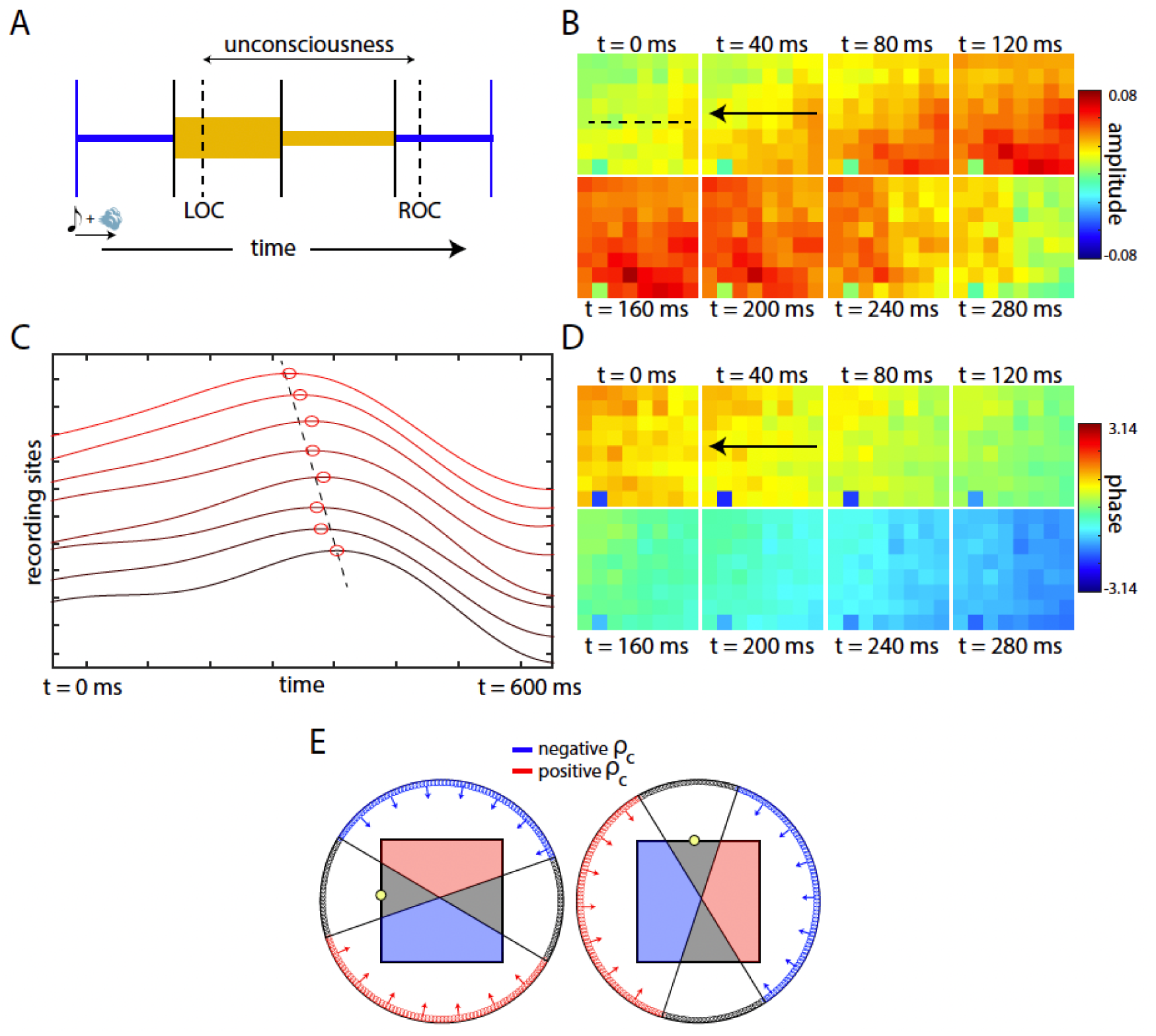
Experimental design and traveling waves. A. Propofol administered in two phases (yellow). The first phase was a higher dose to induce loss-of-consciousness (LOC), followed by a lower dose for LOC maintenance. After propofol cessation recovery of consciousness (ROC) occurred. B. LFP amplitudes (slow-delta) observed across the 8×8 recording array during awake state of Animal 2. Arrow indicates direction of wave propagation. C. Voltage traces from one row of the array (dashed line in B), with peaks marked. Dashed line indicates sequential shifts in peak. D. Phase maps corresponding to amplitude maps in B. E. Wave quantification method based on choice point (yellow). Red region (red arrows) indicates positive coefficient waves, while blue denotes negative. Gray region denotes areas for which wave existence could not be conclusively concluded for that choice point, i.e. the coefficient value was less than the shuffling permutation threshold (see Methods).

Local field potentials were recorded from a total of 6 ‘Utah’ arrays, three in each animal, placed in the ventrolateral prefrontal cortex (vlPFC), the frontal eye fields (FEF), and the parabelt auditory cortex (CPB). Each array consisted of 64 electrodes (8×8 pattern) with a 400-micron spacing. There were 20 experimental sessions (10 for each animal).

### Detecting and quantifying traveling waves

A traveling wave is a sequential activation of adjoining neural groups that gives the appearance of a traveling front of activity. An example is shown in Fig. 1B for an array in the vlPFC. It plots local field potential (LFP) amplitudes band-pass filtered to show the slow-delta band (0.5-3Hz) only. This was recorded during a “baseline” interval before administration of propofol, while the animal was awake. Each tile represents an electrode. Red (blue) indicates higher (lower) LFP amplitudes. Across time, higher amplitudes move sequentially across the array. For example, the first panel (0 ms) shows a high amplitude peak near the right of the array. With each time step, the peak amplitudes “move” towards and through the middle of the array and then towards the left.

Traveling waves could be detected by observing the gradient in oscillation phase values across the recording array (Bhattacharya et al. 2021). Fig. 1C shows oscillatory phases across time on one row of the array (dashed black line in first panel of Fig. 1B). The sequential shift in phase across adjacent electrodes and time is indicative of a traveling wave. The phase plots for the whole array are shown in Fig. 1D. Note the progressive increase in phase from the right to the left. The wave was a not just a “pulse” (a single front or edge of activity moving across the array). A phase gradient could be seen in front of and behind the peak amplitudes, indicating a traveling wave rather than a traveling “pulse”.

We quantified traveling waves using circular-circular correlation coefficients to measure the phase gradient (adjusted for circularity in phase values, see Methods). The coefficient value (*ρ*_*c*_) indicates the spatial correlation between the observed phase map at a given time instant and an idealized rotational phase map around a particular “choice point” on the array. If this correlation was greater than a threshold (determined through a shuffling permutation process, see Methods), a wave was counted for that time instant. A positive vs negative *ρ*_*c*_ indicates wave movement in two opposite directions across the electrode array (Fig. 1E).

To accurately assess wave movement, however, we needed two choice points. This is because the classification of wave directions depended on the choice point. For example, consider the array shown in Fig 1E. Two different choice points on the array are shown in Fig 1E, left and right. With one choice point (circle, Fig 1E, left) waves directed towards the red region in Fig. 1E (left) showed positive correlation (*ρ*_*c*_ >0) while the blue waves showed negative correlation. With a different choice point (Fig. 1E, right), the waves also binned into positive and negative directions but the directions that corresponded to positive vs negative were different. In both cases, some of the waves for one choice point would be directed toward the gray zone (i.e., below threshold for classification as a wave) for the other choice point. Thus, to accurately classify all waves, we needed two points that binned waves in orthogonal directions. This captured all waves with none erroneously falling into the gray zone. If the coefficient for either choice point showed a value greater than the threshold value, a wave was counted. This rendered results independent of any one choice point (Bhattacharya et al. 2021).

### Changes in the number and speed of traveling waves after propofol-induced LOC

We observed traveling waves in three frequency bands (slow-delta = 0.5-3Hz, alpha = 8-12Hz, beta = 12-30Hz). After propofol-induced LOC, there were changes in the number of traveling waves. Fig. 2A shows the wave counts in each of three frequency bands, averaged across sessions and arrays, from the awake state through LOC and then from LOC through recovery of consciousness (ROC). Wave counts in all bands increased shortly after propofol infusion began (before LOC, orange shaded area). The number of alpha (in green) and beta (in blue) waves peaked around the time of LOC. Slow-delta waves numbers (in red) peaked shortly after LOC. Throughout LOC, slow-delta waves showed a large and significant increase in wave counts relative to the baseline period before infusion of propofol (Fig 2A, baseline marked). By contrast, beta waves decreased after LOC, relative to baseline. Alpha waves (8-12Hz) showed an intermediate response, i.e., were more similar to the pre-propofol baseline state than slow-delta and beta. After propofol administration ceased (shown in orange, Fig 2A, right), the wave counts started to move towards pre-propofol baseline values.

**Figure 2:**
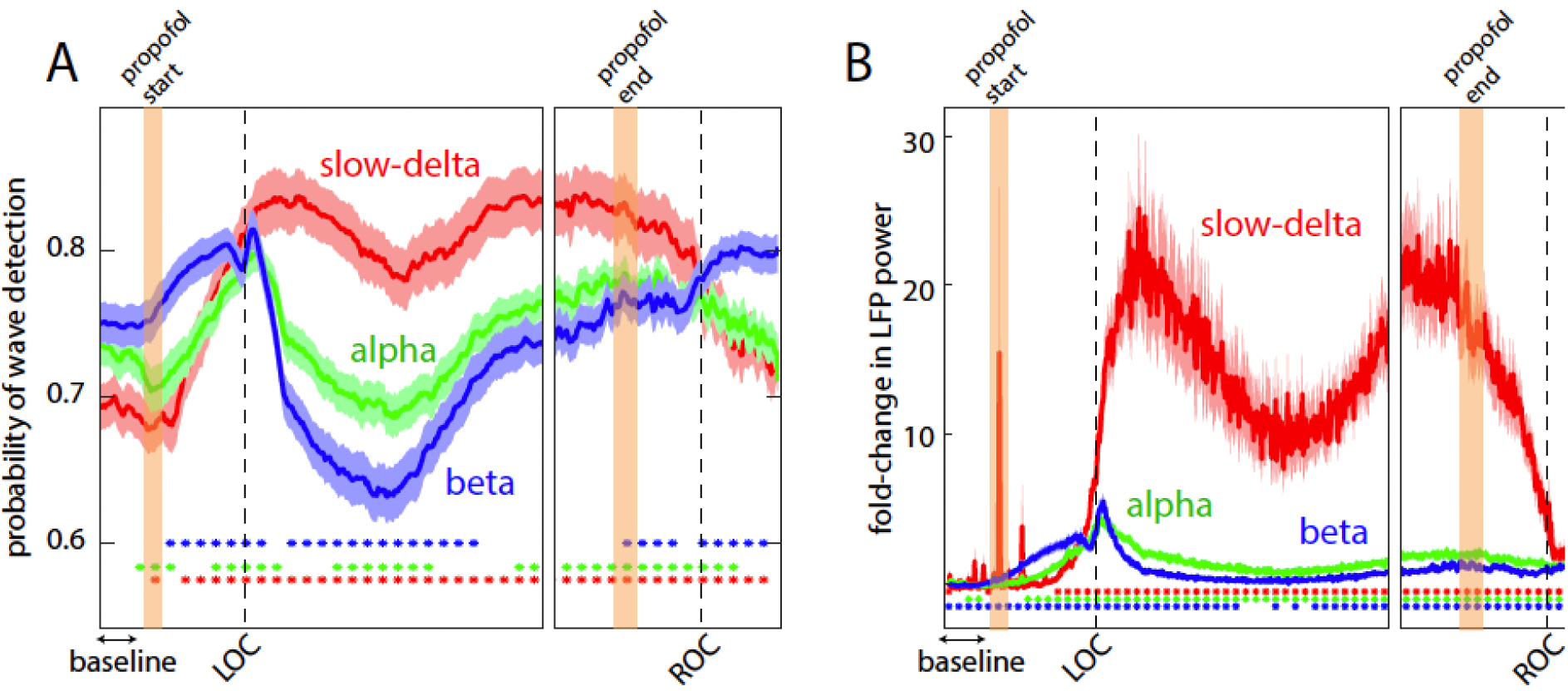
Changes in number of traveling waves and power. A. Probability of wave detection in three frequency ranges across time, averaged across sessions and arrays. The dots denote statistical significance from baseline (marked). B. Fold-change in LFP power compared to baseline (marked) across time, averaged across sessions and arrays. The dots denote statistical significance compared to baseline.

Plots of LFP power (independent of waves, Fig. 2B) showed correspondingly large increases in slow-delta oscillations after LOC relative to alpha and beta. This converged back to baseline levels after propofol cessation.

Propofol also increased traveling wave speed, especially in the slow-delta band. Oscillation frequency and wave speeds, as expected, were positively correlated (Bhattacharya et al. 2021). They were around the 1-20 cm/s range throughout the sessions (Fig. 3A), consistent with other LFP traveling wave studies (Takahashi et al. 2011). The change in traveling wave speeds compared to baseline across time in the session are shown Fig. 3B. Note the large increase in slow-delta wave speed after LOC. Alpha and beta waves also significantly increased in speed but much less so than slow-delta waves. After ROC, wave speeds reduced. They remained significantly higher than the awake state but more similar to their pre-propofolbaseline state.

**Figure 3:**
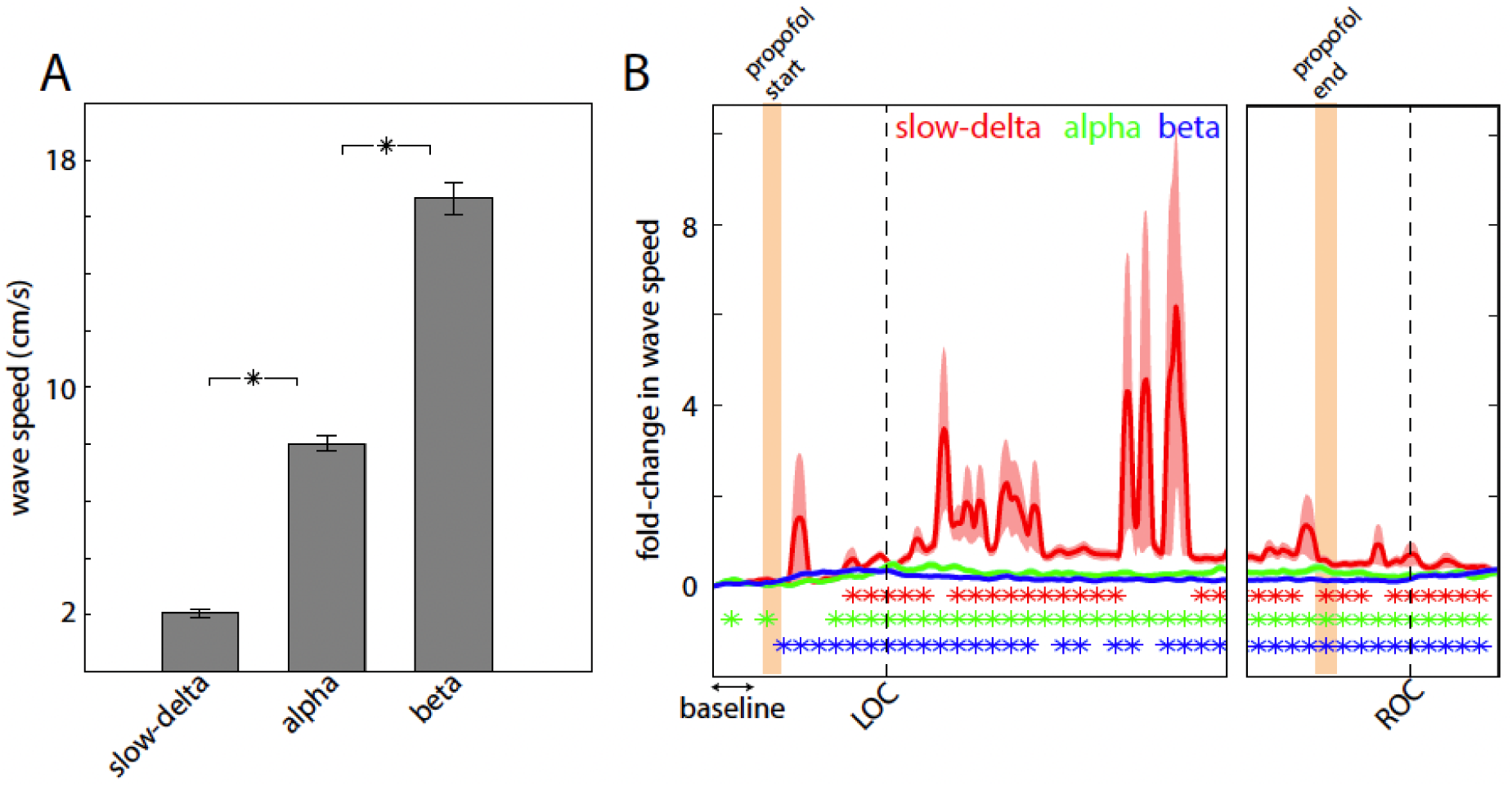
Changes in traveling wave speed. A. Wave speeds observed in the three frequency ranges, averaged across all time (irrespective of propofol), sessions and arrays. Stars denote statistically significant differences. B. Fold-change in wave speeds compared to baseline (marked) across time, averaged across sessions and arrays. The dots denote statistical significance (blue for beta, green for alpha, and red for slow-delta compared to baseline).

### Changes in the spatial structure of traveling waves

Propofol also changed the spatial organization of the waves. Specifically, one noticeable change was in their spatial coherence, i.e., whether they were more “solid” waves or “broken” waves. A solid wave would show similar amplitudes across adjacent electrodes and smooth continuous changes across progressively further electrodes. In other words, they had a uniformly organized structure. A broken wave would, by contrast, show a larger range of amplitude values on the array. This is illustrated for simulated data in Fig 4A. The top shows a more solid wave and the bottom a broken wave. We quantified this using the coefficient of variation (COV: standard deviation divided by the mean) of the amplitude envelope of the traveling wave. Simulated broken waves showed a significantly higher amplitude COV when compared to the more spatially coherent wave (Fig. 4B).

**Figure 4:**
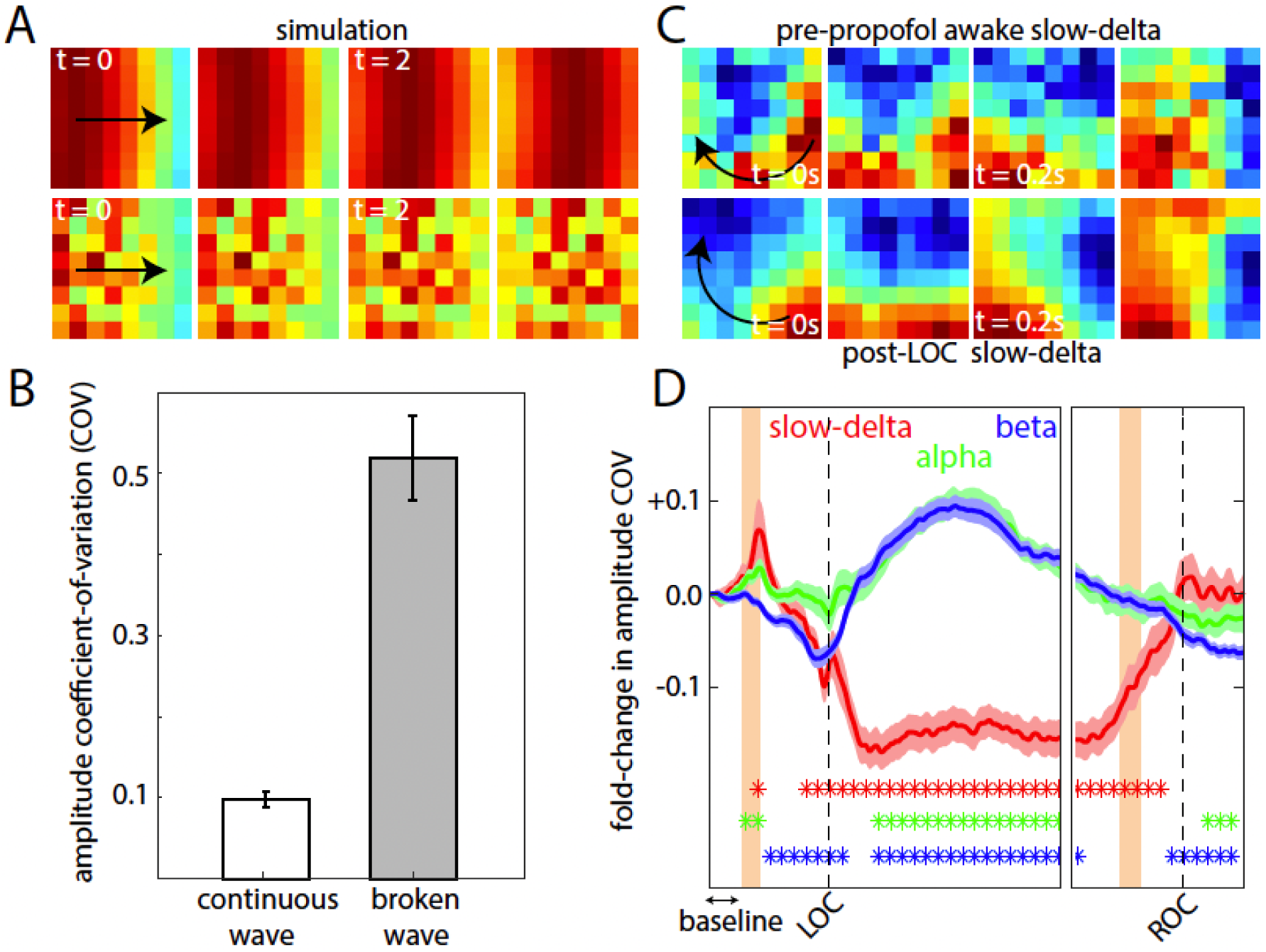
Changes in traveling wave structure. A. Simulations of traveling waves with different levels of spatial couplings across adjacent elements. Top, continuous smooth wave with homogenous amplitude map, and bottom, broken wave with heterogenous amplitude map across the array. B. Quantification of the coefficient of variation (COV) of amplitudes observed across the array for the simulated continuous and broken waves. C. Example of slow-delta waves observed in the pre-propofol state (top) and unconscious state (bottom) for Animal 1. D. Quantification of the fold-change in amplitude COV for slow-delta, alpha, and beta waves compared to baseline (marked) averaged across sessions and arrays. The dots denote statistical significance (blue for beta, green for alpha, and red for slow-delta compared to baseline).

We found that slow-delta traveling waves increased their spatial coherence (became more solid) after LOC, while alpha and beta waves did the opposite. Fig. 4C shows examples of slow-delta waves (moving in the direction of the black arrow) on the vlPFC recording array of Subject 1, in the pre-propofol baseline state (top) and after LOC (bottom). The post-LOC slow-delta wave had more amplitude homogeneity than its baseline counterpart indicating that it was more solid. The change in COV (compared to pre-propofol baseline) for slow-delta and beta waves averaged across all arrays and sessions is shown in Fig. 4D. Slow-delta waves showed significantly lower amplitude COV after LOC, indicating more spatial coherence. By contrast, alpha and beta waves became more broken and less structured after LOC, i.e. they showed a significant increase in COV relative to the baseline. After ROC, slow-delta waves returned to their baseline coherence. Beta (and to some extent alpha) waves not only regained their structure post-ROC but showed a “rebound”. They showed even stronger spatial coherence (lower amplitude COV) compared to their pre-propofol baseline state.

### Propofol changed traveling wave patterns

In a previous study (Bhattacharya et al. 2021), we demonstrated how one can leverage the properties of three circular-circular correlation coefficients to distinguish between rotating and planar waves (Fig. 5A). We applied the same methods here (see Methods). Both rotating and planar waves were observed in all the recording arrays. On average, during the pre-propofol baseline, planar and rotating slow-delta waves had similar incidence on the arrays (Fig 5B).

**Figure 5:**
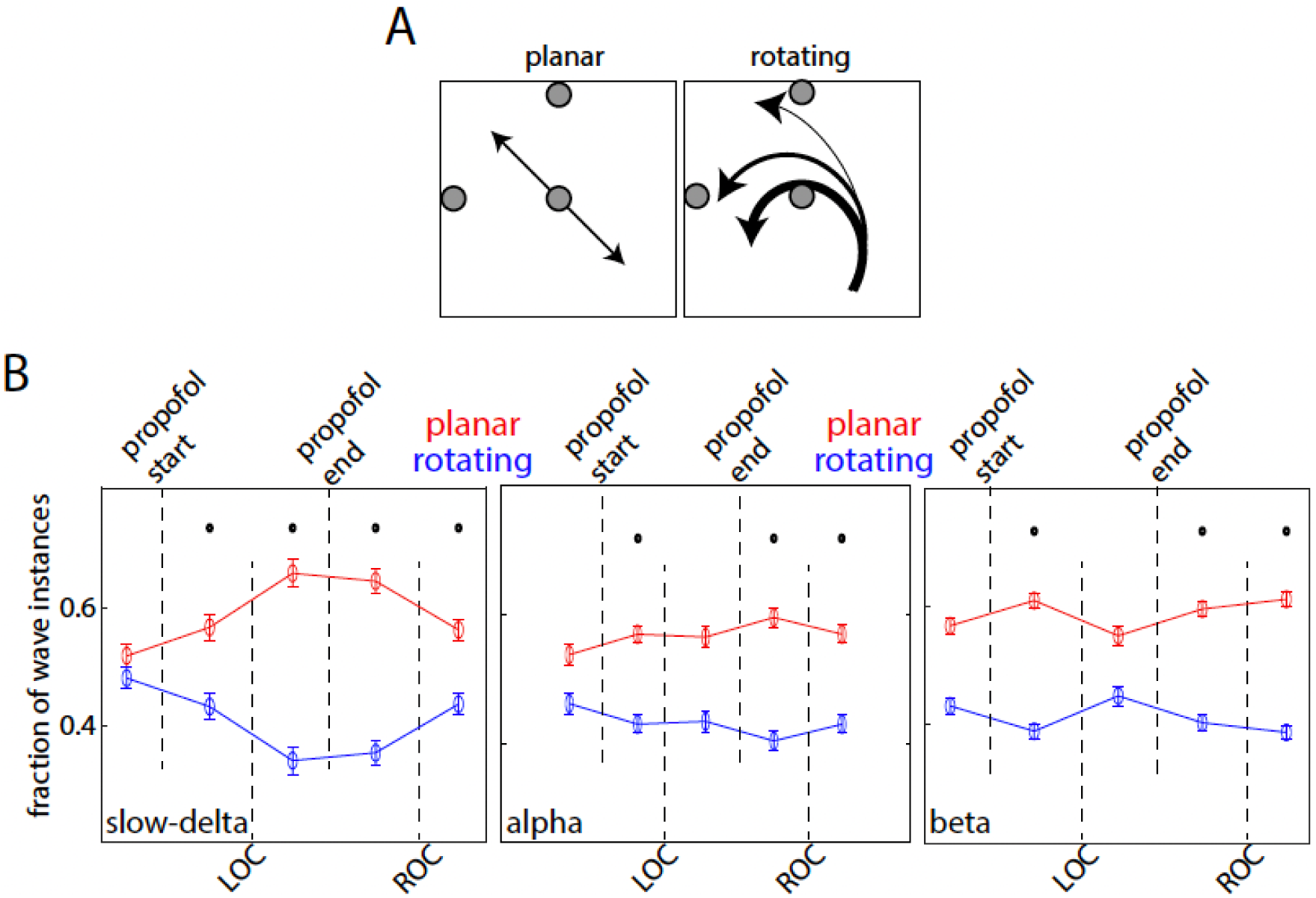
Changes in traveling wave pattern. A. Planar and rotating wave examples, detected with three choice points on the array (grey circles). B. Fraction of planar and rotating waves noted at different stages of anesthesia for slow-delta (left), alpha (middle), and beta (right) waves averaged across sessions and arrays. Dots denote statistical significance from baseline.

After LOC, there was a significant decrease in slow-delta rotating waves and increase in slow-delta planar waves (Fig 5B). This distinction was evident after propofol induction and persisted through LOC. After ROC, slow-delta planar and rotating waves started to converge to their baseline values. By contrast, the changes in alpha and beta wave patterns were more modest. Alpha waves showed the same trends as slow-delta waves. Planar waves increased and rotating waves decreased, albeit to a lesser extent than the slow-delta waves. Upon propofol induction, there was a decrease in beta rotating waves and a modest but significant increase in planar waves (Fig. 5B). Unlike slow-delta, the incidence of beta planar vs rotating waves post LOC was similar to that during baseline. The changes reappeared after propofol cessation and in ROC but were modest as well. It is worthwhile to note here that the arrays typically captured only part of the rotating waves. Thus, rotating waves could appear to become planar if the rotating waves shifted in anatomical location. Nonetheless, these results show that propofol induced changes in wave patterns, especially in the slow-delta band.

### Propofol made slow-delta traveling waves more stereotyped

Two waves are similar when they not only have the same direction, but also have similar speed, spatial coherence, and overall organization. They can also be “anti-similar”, i.e., they are similar in other properties but flow in opposite directions (like a mirror-image). We found that after propofol-induced LOC, there was an increase in slow-delta wave anti-similarity.

To quantify this, we performed a similarity analysis on the wave patterns on each array. We calculated, for each array separately, the spatial correlation between each wave at a given snapshot in time with all other waves at other snapshots in time. A positive correlation indicates repetition of a particular wave pattern (i.e., high similarity across time). A negative correlation indicates “anti-similarity”, i.e. repetition of waves that had high similarity across time but moved in opposite directions. Correlations around zero indicate randomness in wave properties.

Propofol caused an increase in slow-delta wave anti-similarity, relative to baseline. In other words, slow-delta waves became more stereotyped. Fig. 6A shows the wave similarity values averaged across arrays and sessions. The distribution for slow-delta traveling waves changed from a more unimodal histogram centered around zero similarity during baseline to a bimodal histogram after LOC. This meant that slow-delta wave properties were relatively random during baseline but, after LOC, shifted to a repetitive pattern of mirror-image similar waves flowing in opposite directions. This was evident in both early and late periods of unconsciousness (post-LOC) (Fig. 6A). Then, after ROC, slow-delta waves returned to a unimodal random organization similar to baseline. By contrast, alpha and beta waves (Fig. 6B, C) did not show any shift in the distribution of wave similarity. They were relatively random during baseline and remained so after LOC.

**Figure 6:**
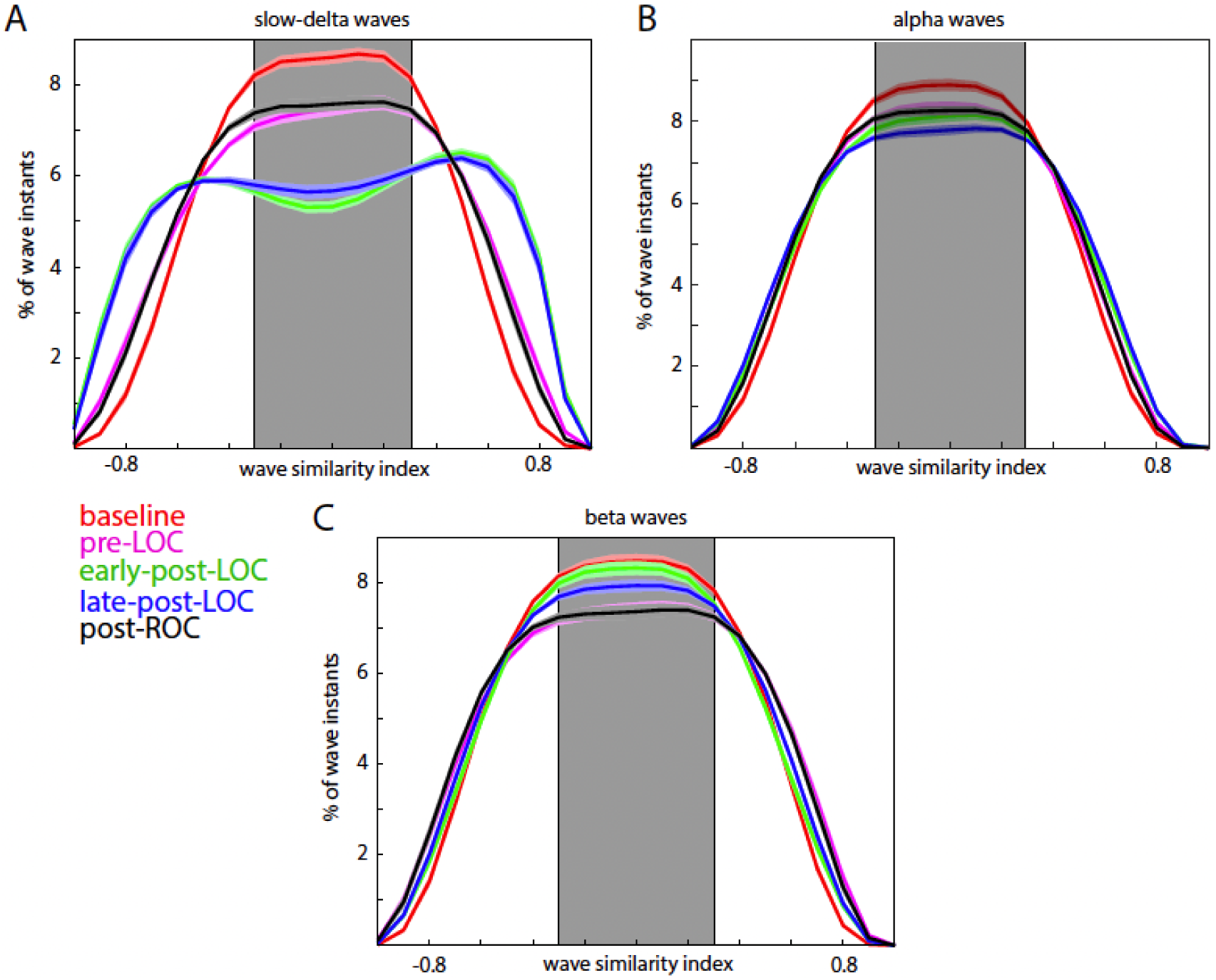
Slow-delta traveling waves become more stereotyped. Similarity indices noted for slow-delta (A), alpha (B), and beta (C) waves at different stages of anesthesia, averaged across sessions and arrays. Pre-LOC denotes the period where propofol administration has started but the subject is conscious. Shaded region denotes low similarity values (threshold determined by shuffling permutation procedure, see Methods).

### Propofol caused slow-delta and higher-frequency waves to flow in mutually exclusive directions

Traveling wave directions were also altered after LOC. To quantify this, we classified traveling waves into four different directions. As before, we used two circular-circular correlation coefficients from two choice points in order to capture all wave directions. They were then binned into 4 directions – towards the four quadrants shown in Fig. 7A.

**Figure 7:**
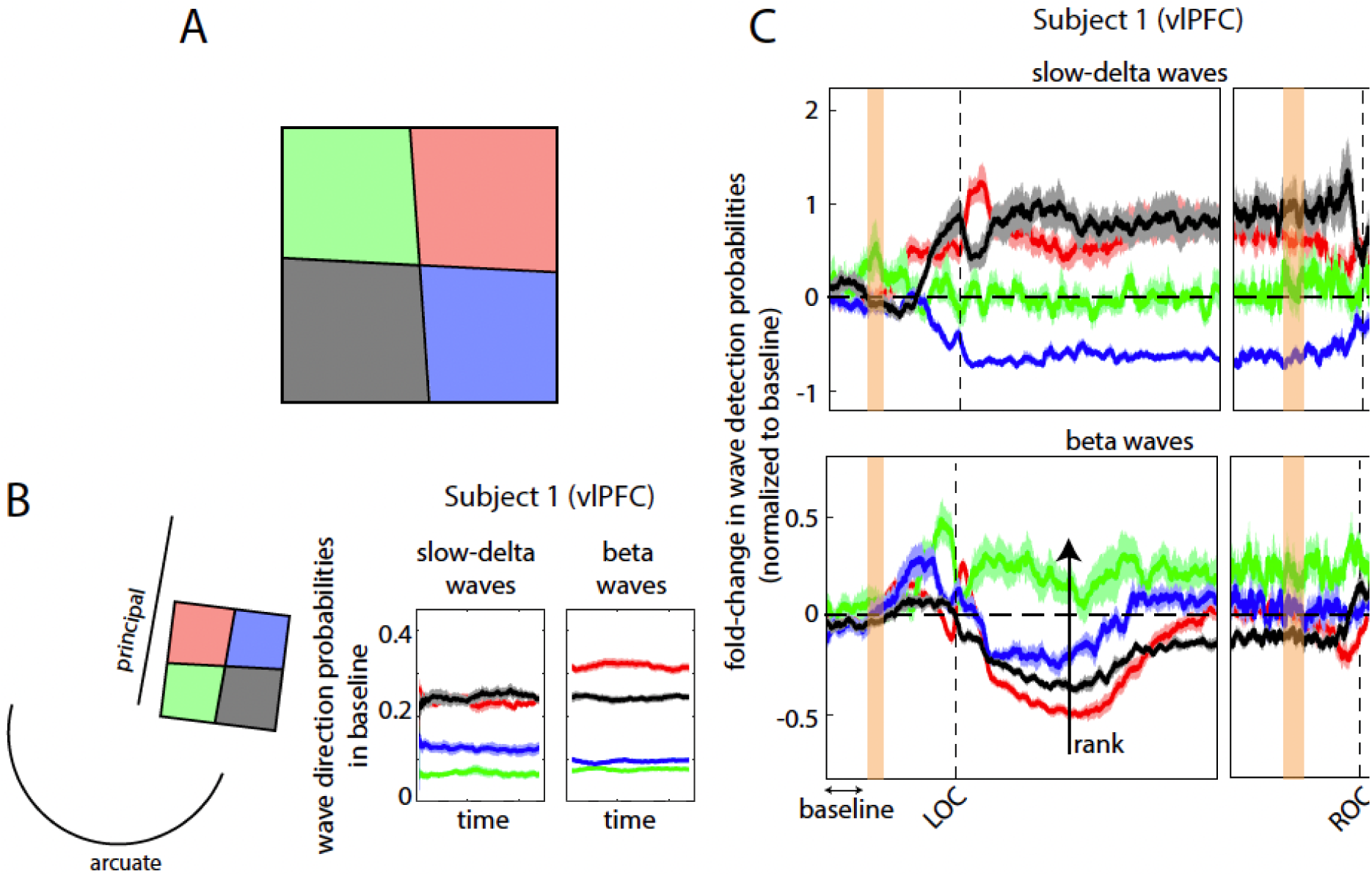
Traveling waves change direction. A. Four direction bins (red, green, blue and black) along which waves were categorized. B. Location of vlPFC array of Animal 1 relative to brain sulci (left), and baseline levels of wave direction probabilities (right) for the same array. C. Fold-change in wave detection probabilities in each of the four directions compared to baseline, across time for the same array in B. Black arrow in beta panel indicates how wave directions were ranked in this particular case (green got rank 1, while red got rank 4).

An example of wave-direction analyses is shown in Fig. 7B. It plots the probability of detecting waves in the four binned directions for slow-delta and beta waves during baseline for the vlPFC array of Subject 1 (array location relative to brain sulci is shown on the left). Both slow-delta and beta waves flowed more frequently toward the “red” and “black” zones (see Fig 7A), that is, toward “upper right” and “lower left” of the array. Note that waves from different frequency bands did not necessarily flow in the same directions, as seen here. On other arrays, waves from different frequency bands could have different preferred directions.

Fig. 7C shows the change in wave direction on the vlPFC array of Subject 1 post-LOC. Fold-change was calculated by dividing the probability of wave detection in each of the four directions with the probability of wave detection in the same direction during baseline. Fig. 7C shows this plotted across time averaged across sessions. Slow-delta waves increased towards the directions preferred during baseline (i.e., the red and black directions). By contrast, beta waves reduced in these directions. Instead, they increased towards the green section (“upper left”). Notably, the green direction was least preferred for beta waves during baseline (Fig. 7B right). Post-ROC the wave directions returned to their baseline preferences.

This segregation of slow-delta and beta waves into different directions post-LOC was seen across all arrays. The six arrays were each oriented differently with respect to brain sulci. Plus, the waves were oriented differently with respect to the arrays (Bhattacharya et al. 2021). Thus, we could not simply combine the same directions across arrays (e.g., red to red, black to black). Instead, we rank-ordered the directions based on which directions changed the most from baseline to LOC for beta (black arrow Fig. 7C). That is, the direction that saw the most increase in beta waves after LOC was rank 1, and the direction that saw the least increase (or most decrease) was rank 4. We applied the same rank ordering from beta to alpha and slow-delta in order to compare how the directions changed relative to one another. Directions with the same rank were combined across arrays (color coded: red:1, green:2, blue:3, black:4).

Fig. 8 shows this analysis averaged across all arrays. Post-LOC slow-delta and beta (and alpha) waves increased in mutually exclusive directions. Beta waves increased in one direction (rank 1, red, by definition) after LOC. By contrast, there was a decrease in beta waves in the other directions (green, blue, and black). Alpha waves behaved similarly as beta waves. Slow-delta waves showed opposite trends to that of beta waves. They increased in the green direction especially but also in the black and blue directions. The red direction, which increased in beta waves, showed a decrease in slow-delta waves. After ROC, wave direction preferences converged (or started to converge) to their pre-propofol baseline levels (Fig. 8).

**Figure 8:**
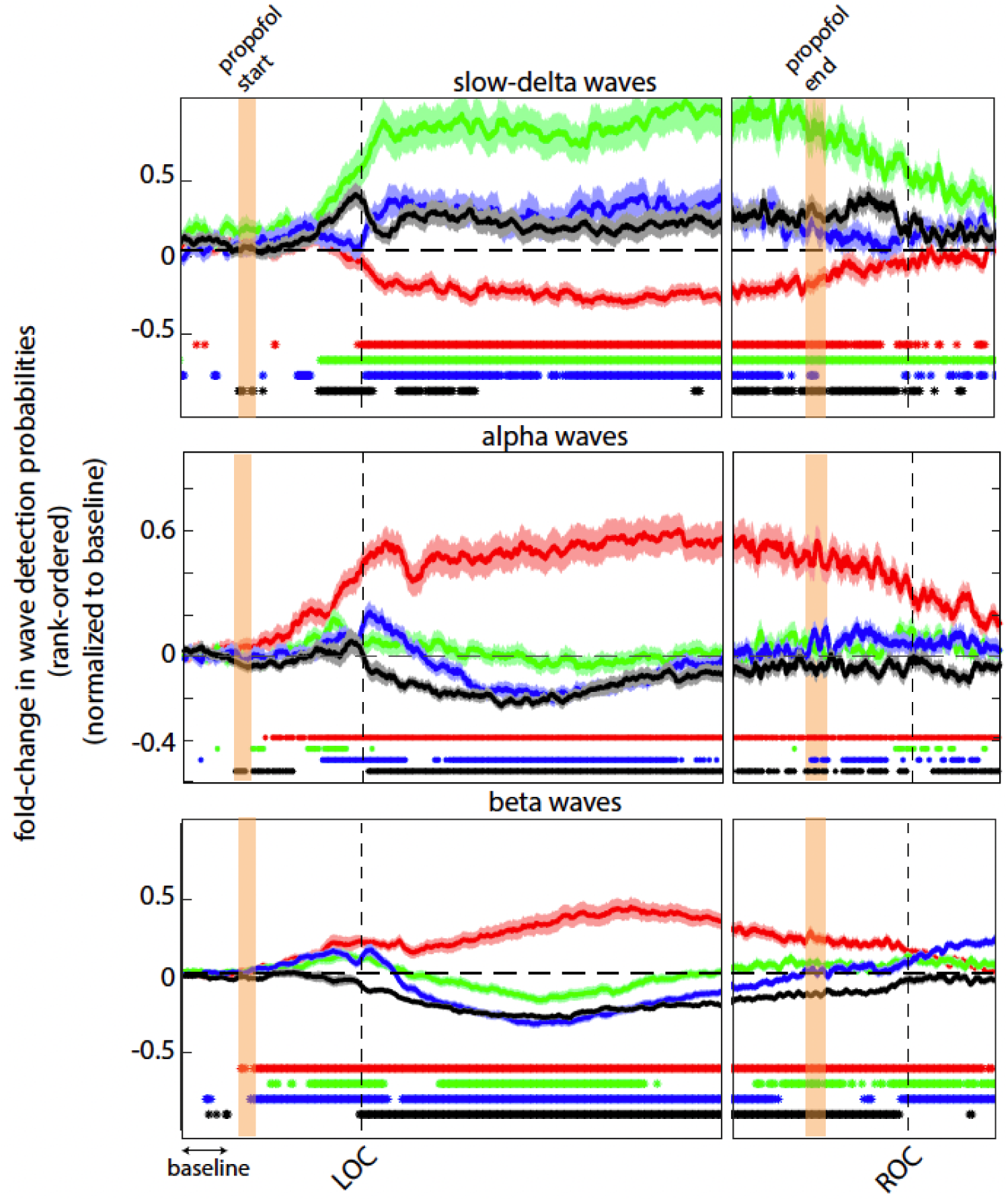
Traveling waves in different bands flow in different directions. Fold change in wave detection probabilities averaged across all arrays – combined through a rank-ordering system (ranked according to beta wave-change post-LOC as shown in Fig. 7). Each color indicates a particular wave direction. Dots denote statistical significance for each direction compared to baseline.

## Discussion

We found that after propofol-induced loss of consciousness (LOC), cortical traveling waves were altered. Slow-frequency delta (∼1 Hz) waves increased while higher-frequency (8-30 Hz) waves decreased. The slow-delta waves sped up and became more spatially organized. They became more planar (and less rotating) and increased mirror-image waves traveling in opposite directions. Whatever directions slow-delta waves flowed in after LOC, they dominated. Higher-frequency waves decreased and lost structure after LOC, despite showing increased LFP power, and flowed preferentially in directions where slow-delta waves were less frequent.

This is consistent with prior work showing increases in slow frequency power and coherence in cortex following propofol-induced LOC (Bastos et al. 2021; Redinbaugh et al. 2020; 2021). Our results are also consistent with prior observations that traveling waves (under anesthesia) have preferred directions and mirror-reflective properties (Sanchez-Vives and McCormick 2000; Mitra et al. 2018; Xu et al. 2007). They also support a hypothesis by Muller et al (2018) (Muller et al. 2018). They posited that decreased spiking activity under anesthesia, as seen in our earlier work (Bastos et al. 2021), can lead to a greater recruitment of neuron groups into a traveling wave. The result of anesthesia would be more “solid”, organized, waves than sparse “broken” waves, which is what we observed. Traveling waves under anesthesia also seem to cross anatomical boundaries in the visual cortex while those in awake state do not (Muller et al. 2014; Xu et al. 2007). This could be explained by our observation of increased slow-delta wave speed and organization after LOC. It could allow waves to travel longer distances without loss of structure (Bhattacharya and Iglesias 2019). Plus, we observed a decrease in rotating, and an increase in planar, slow-delta waves after LOC. Planar waves can traverse larger distances, as rotating waves tend to lose structure away from their core (Bhattacharya et al. 2021). Our observations were consistent across the frontal and auditory cortices of NHPs. However, a study in the somatosensory cortex of freely moving mice showed that stimulation-induced traveling waves spread farther when compared to those in anesthetized mice (Ferezou, Bolea, and Petersen 2006). The differences between this study and ours could be due to use of a different anesthetic, different species, or a different cortical area (somatosensory cortex). Reimer et al. (2011) (Reimer et al. 2011) reported that traveling waves in rat cortex did not change significantly under different anesthetics (nitrous oxide, isoflurane or ketamine) but they did not test propofol. However, we hypothesize that our results may extend to other anesthetics as well specifically because isoflurane and propofol have similar mechanisms of action through GABAergic circuits (Franks and Lieb 1994).

The decrease in rotating waves might also help explain differences between sleep and anesthesia. Rotating waves have been associated with memory consolidation during sleep spindles (Muller et al. 2016). It has been hypothesized that rotating waves can precisely control spike timing relationships. This could foster the spike-time-dependent plasticity that consolidates memories during sleep (Muller et al. 2016). Given the link between rotating waves and sleep, the “un-rotating” of traveling waves by propofol may disrupt the spike-time-dependent plasticity needed for memory consolidation and could potentially explain anesthesia-induced retrograde amnesia.

Unlike slow-delta waves, beta and alpha waves decreased in number and lost structure. It was as if the slow-delta waves were “swamping” cortex and crowding out the higher frequency traveling waves. Beta and alpha waves did not disappear, though. They seemed to be redirected by the slow-delta waves. After LOC, higher frequency (8-30 Hz) waves reoriented to a direction not being used by slow-delta waves. We can perhaps think of wave directions as specific neural pathways/channels. Slow-delta waves after LOC became stronger, more frequent, and more directional. This “crowded” the pathways where slow-delta waves flowed but freed up other pathways to which the higher frequency waves “retreated”, often using channels that were low-preference directions in the awake state.

We observed that both slow-delta (0.5-3 Hz) and beta (12-30 Hz) oscillatory power increased after propofol-induced LOC – albeit the former significantly more than the latter. Beta rhythms are thought to support cognitive functions in NHPs (Bastos et al. 2018; Lundqvist et al. 2018). Our results indicate that the spatial structure of these rhythms may be important markers of cognition as well. While beta power increases somewhat under propofol, beta traveling waves decreased, lost structure, and seemed to be forced to change directions by slow-delta waves. This suggests a mechanism by which propofol may cause unconsciousness: slow-delta waves dominate and disrupt the traveling waves in the higher frequencies associated with cognition.

## Materials and Methods

### Subjects, LFP recordings and propofol administration

Experimental data was used from our earlier paper (Bastos et al. 2021). Two rhesus macaques (*Macaca mulatta*) aged 14 years (Subject 1, male,∼13.0 kg), and 8 years (Subject 2, female,∼6.6 kg) participated in these experiments. Detailed surgical and housing protocols can be found in (Bastos et al. 2021).

Experimental sessions were carried out in two phases. In the first phase, a period of 15–90 min of awake baseline activity was recorded. Next, propofol was intravenously infused via a computer-controlled syringe pump (PHD ULTRA 4400, Harvard Apparatus, Holliston, MA). The infusion protocol was stepped such that unconsciousness was induced via a higher rate infusion (285 mcg/kg/min for monkey 1; 580 mcg/kg/min for monkey 2) for 20 min before dropping to a maintenance dose (142.5 mcg/kg/min for monkey 1; 320 mcg/kg/min for monkey 2) for an additional 40 min.

Facial movements and pupil size were tracked by infrared (Eyelink 1000 Plus, SR-Research, Ontario, CA) throughout the sessions. The instant of eyes-closing that persisted for the remainder of the infusion was marked as loss of consciousness (LOC). ROC was marked by the instant of the first to occur between eyes reopening or regaining of motor activity following propofol infusion cessation. Further details can be found in (Bastos et al. 2021). All procedures followed the guidelines of the MIT Animal Care and Use Committee (protocol number 0619-035-22) and the US National Institutes of Health.

The subjects were chronically implanted with 8×8 iridium-oxide “Utah” microelectrode arrays (1.0 mm length, 400 μm spacing; Blackrock Microsystems, Salt Lake City, UT) in the ventrolateral prefrontal cortex (vlPFC), frontal eye fields (FEF) and the auditory parabelt cortex (CPB). Signals were recorded on a Blackrock Cerebus. LFPs were recorded at 30 kHz and filtered online via a lowpass 250 Hz software filter and downsampled to 1 kHz. All preprocessing and analysis were performed in Python or MATLAB (The Mathworks, Inc, Natick, MA). For the power analysis, the resulting signals were convolved with a set of complex Morlet wavelets.

### LFP spatial phase maps

The raw LFP traces were filtered in the desired frequency range, using a 4^th^ order Butterworth filter for alpha (8-12 Hz) and beta (12-30 Hz) ranges, and a 3^rd^ order Butterworth filter for slow-delta (0.5-3Hz) oscillations, forward-reverse in time to prevent phase distortion (see MATLAB function *filtfilt*). A Hilbert transform was used to obtain the analytical signal for each electrode. The phase of each electrode for the 8×8 array is called the “phase map” for that time instant. These phase maps (unsmoothed) were checked for gradients to identify traveling waves.

### Shuffling procedure

To ensure that the probability of detecting traveling waves exceeded that expected by chance – we performed a random shuffling procedure to establish a threshold for the correlation coefficient – beyond which a traveling wave was counted. This was done by shuffling the phase values on the array randomly (with 25 different types of random permutations) and calculating the correlation coefficient. The 99^th^ percentile of the resulting distribution of coefficient values determined a threshold (0.3) above which the correlation exceeded chance.

### Traveling wave identification and classification

We used circular statistics to identify wave patterns. Methods followed our previous investigation of traveling waves in the prefrontal cortex (Bhattacharya et al. 2021).

The circular-circular correlation coefficient reports the spatial gradient similarity between two phase maps that are adjusted to account for circular phase values. For the signal phase (*φ* at each array coordinate [a,b]) and the rotation angle (*θ*) around the chosen point, the circular-circular correlation coefficient thus was:

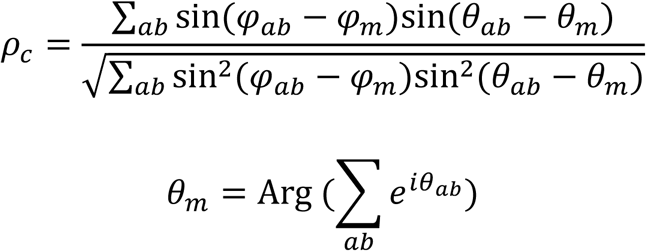

As we showed in our previous study (Bhattacharya et al. 2021), the choice of point around which the coefficient was calculated split the array into two regions (Fig. 1E). A traveling wave towards the positive half (red arrows, Fig. 1E) would result in positive values, while the opposite direction would result in negative values. Each chosen point had a “chance zone” around which the coefficient values would be too low (less than the threshold determined by the shuffling permutation procedure) to make a conclusion regarding wave existence. Hence two points, (1,4) and (4,1), chosen such that their chance zones did not overlap, were used to determine wave existence (Fig. 1E). If either point reported a coefficient value greater than the permutation threshold, a wave was counted. Further, a combination of the two coefficient values could thus be used to bin the waves into four directions (Fig. 7A): *ρ*_*c*14_>0, *ρ*_*c*41_>0 (red direction), *ρ*_*c*14_>0, *ρ*_*c*41_<0 (green direction), *ρ*_*c*14_<0, *ρ*_*c*41_>0 (blue direction), and *ρ*_*c*14_<0, *ρ*_*c*41_<0 (black direction). As this was an 8×8 array (even numbers) it is understandable that the bisecting axis was not perfectly horizontal for *ρ*_*c*14_ or perfectly vertical for *ρ*_*c*41_. Also, it is important to note that our methods were not dependent upon the exact choice of points (Bhattacharya et al. 2021).

To distinguish between planar and rotating waves, exactly similar to our earlier study (Bhattacharya et al. 2021), we used a third rotation map around (4,4) along with the (1,4) and (4,1) maps. Each wave instant thus had its associated three coefficient values: *ρ*_*c*14_, *ρ*_*c*41_ and *ρ*_*c*44_. Using simulations, we obtained similar coefficient values for different types of planar and rotating waves. Waves were simulated using the following equations (Muller et al. 2016):

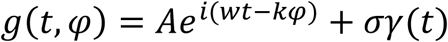

where *φ* was the input phase map, *w* was the temporal frequency and *k* was the spatial wavenumber (1/wavelength). The second term was a Gaussian white noise term, with zero mean and standard deviation *σ*.

In this way, we obtained the three coefficient values for simulated waves, to go with our experimental coefficient dataset. We compared these values to automatically classify the type of wave observed, based on the Euclidean distance between a wave type and the observed phase map. Coefficient matching with three different maps, allowed for greater accuracy with lesser chances of misclassification.

Wave speed at a time instant was calculated from the phases (*p*) by dividing the temporal frequency (*∂p*/*∂t*) at that time with the spatial frequency (*∂p*/*∂∂*) (H. Zhang and Jacobs 2015). The gradients obtained were averaged across electrodes to get the net wave speed for that time instant.

### Wave spatial coherence

The amplitude envelope obtained from the Hilbert transform of the LFP signal was used to determine the spatial coherence of the traveling wave. A wave was deemed to be spatially coherent when it showed similar amplitudes across the array at a particular instant. A broken wave would show a larger amplitude variance across electrodes on the same array. We demonstrated this using two simulations – one, where the array elements had uniform oscillation amplitude ranges, and two, where the amplitude ranges were randomly distributed (analogous to a fragmented cortex). When the same phase gradient (traveling wave, left to right, Fig. 3A) was imposed in both cases, the first showed a smooth, organized wave structure (Fig. 3A, top), while the other showed a broken traveling wave with heterogeneous amplitudes across the wave band. For the broken wave, the amplitude envelope showed larger variance when compared to the more solid wave. Coefficient of variation (COV) was defined as the standard deviation divided by the mean.

### Wave similarity analysis

Two phase maps were checked for similarity by computing the circular-circular correlation between them (Muller et al. 2016). A high positive coefficient value indicated similar waves, i.e. waves with the same phase organization on the array. A high negative coefficient value indicated anti-similar mirror-image type waves. A coefficient around zero (shaded region, Fig. 6, with the threshold determined by the shuffling permutation procedure) indicated no conclusive similarity between the phase maps. This analysis was done across all time instants in each session.

## Acknowledgments

We thank Jesus Ballesteros, Andre Bastos, Alex Major, Dimitris Pinotsis, and Jefferson Roy for helpful comments. These studies were supported by NIMH R01MH11559, NIGMS P01GM118269, and the MIT Picower Institute Innovation Fund.

## Author Contributions

EKM and ENB designed and supervised the experiments. JAD, SLB and MM performed the experiments. SB conceived and performed the analysis. SB, ENB and EKM wrote the manuscript.

